# Unique mushroom spine morphology signature in cocaine relapse ensembles in rat nucleus accumbens core

**DOI:** 10.64898/2026.07.15.738740

**Authors:** F. Javier Rubio, Emem Ukpong, Rajtarun Madangopal, Daniel E. Olivares, Elias M. Hilaire, Akeem Henry, Amaya Kelly, Louisa Kane, Carlos Mejias-Aponte, Bruce T. Hope

**Affiliations:** Neuronal Ensembles in Addiction Section, Behavioral Neuroscience Research Branch, National Institute on Drug Abuse IRP, NIH, Baltimore, MD; Laboratory of Sensorimotor Research, National Eye Institute, NIH, Bethesda, MD

**Keywords:** ensemble, engram, operant learning, relapse, novel context, dendritic spines, interspine intervals, clustering, mushroom spines

## Abstract

Fos-expressing neuronal ensembles in nucleus accumbens (NAc) are selectively activated by drug-associated cues and shown to play a causal role in drug-cue memories. Many unique molecular and cellular alterations have been identified in these ensembles and thought to be components of an engram encoding the memory; however, spine morphology alterations on these ensembles have rarely been examined. Here, we examined alterations of identified spine types on Fos-expressing neurons in NAc core that were selectively activated by drug-associated cues during cocaine relapse. We combined Fos immunolabeling along with viral-based sparse GFP labeling to identify Fos-positive and Fos-negative neurons with distinguishable dendrites and quantified spine composition, densities and interspine intervals (ISIs) for all spine types, and head and neck diameters of mushroom spines. ISIs and head and neck diameters of mushroom spines were altered on Fos-positive (versus Fos-negative) neurons activated during cue-induced cocaine relapse. Critically, these alterations were not found on Fos-positive neurons activated by exposure to a novel context on relapse test day, despite having identical cocaine self-administration histories. Based on previous work, this suggests these alterations were not simply due to acute activation of a randomly selected set of neurons on test day, but rather they were altered specifically on neurons selectively activated during cocaine self-administration training and then reactivated during relapse. Stubby spines and immature spines, including filopodia and thin spines, were not altered. Altogether, these mushroom spine alterations describe a unique spine morphology signature for a drug-cue memory in NAc neuronal ensembles selectively activated during cocaine relapse.

## Introduction

During repeated experiences with drugs of abuse, people and animals learn to associate a variety of cues in the drug environment with the rewarding effects of the drug that can then trigger memories of drug reward weeks to years later and promote drug relapse [1]. These drug-cue memories are thought to be encoded within specific patterns of sparsely distributed neurons called neuronal ensembles that are selectively activated by the drug-associated cues [2–4]. The neuronal ensembles are commonly identified by expression of immediate early genes (IEGs) such as Fos, Arc or Npas4 during memory reactivation and selective manipulations of these ensembles in nucleus accumbens (NAc) and other mesolimbic forebrain areas demonstrate that they play a causal role in encoding drug-cue memories.

Unique cue-induced molecular and cellular alterations are induced within these ensembles and thought to contribute to long-lasting physical traces or engrams encoding the drug-cue memories [5]. Dendritic spine alterations specifically on Fos-expressing ensemble neurons in NAc can play a similar role in these engrams but they have rarely been examined. The principal neurons in NAc and its neuronal ensembles are medium spiny neurons (MSNs) with 10,000 to 30,000 dendritic spines on each neuron [6]. Spines on distal dendrites receive glutamatergic terminals carrying information about drugs and cues from cortical, amygdala, and other brain areas while their proximal dendrites receive input primarily from local interneurons [7]. These spines are highly dynamic and can mature from filopodia into thin, stubby, and ultimately stable mushroom spines that are associated with long-term memory storage [8–13]. Activity-dependent remodeling of mushroom spines, in particular, is considered a key component of synaptic plasticity underlying the formation of engrams encoding long-term memories [14–19].

While previous studies assessed dendritic spines in NAc and other brain areas following drug learning [20–22], very few have examined spine alterations specifically on Fos-expressing ensemble neurons [23,24]. In the current study, we examined dendritic spine alterations on Fos-positive neurons versus Fos-negative neurons in NAc core following an operant model of cue-induced cocaine relapse [25]. We assessed spine types and densities, and interspine intervals (ISIs) for each of the different morphological spine types in both distal and proximal dendrites, and mushroom spine head and neck diameters, on NAc neuronal ensembles previously shown to encode cocaine relapse in rats [26]. Critically, we compared dendritic spine alterations on Fos-expressing neurons activated by cue-induced cocaine relapse versus alterations on a different set of Fos-expressing neurons activated by a distinct set of cues in a novel context following the same cocaine self-administration training.

## Results

### Cocaine self-administration training and seeking test

The experimental design is shown in Fig. 1A. Rats were trained to self-administer cocaine by pressing a lever paired with a cue light for 12 days followed by 21-28 days of forced abstinence. Rats demonstrated reliable cocaine self-administration as indicated by a significant increase in the number of infusions (t _(1,43)_ = 13.8, p < 0.01) and active lever presses (t _(1,43)_ = 11.3, p < 0.01) when lowering the cocaine dose from 0.75 mg/kg per infusion on training days 1-6 to 0.375 mg/kg per infusion on training days 7-12 (Fig. 1B). After training, rats were returned to their homecages for 3-4 weeks of experimenter-imposed abstinence. On test day, rats were separated into three groups and exposed to either (1) the Training context previously associated with reward (Training context) or (2) a Novel context or (3) were kept in their home cages on test day (No Test group); all groups were previously matched for similar numbers of infusions and active lever presses during training. Only the Training context group underwent cue-induced seeking on test day; non-reinforced responding on the previously active cocaine-paired lever was significantly greater than inactive lever presses (t _(1,14)_ = 8.1, p < 0.01) (Fig. 1C).

**Figure 1.**
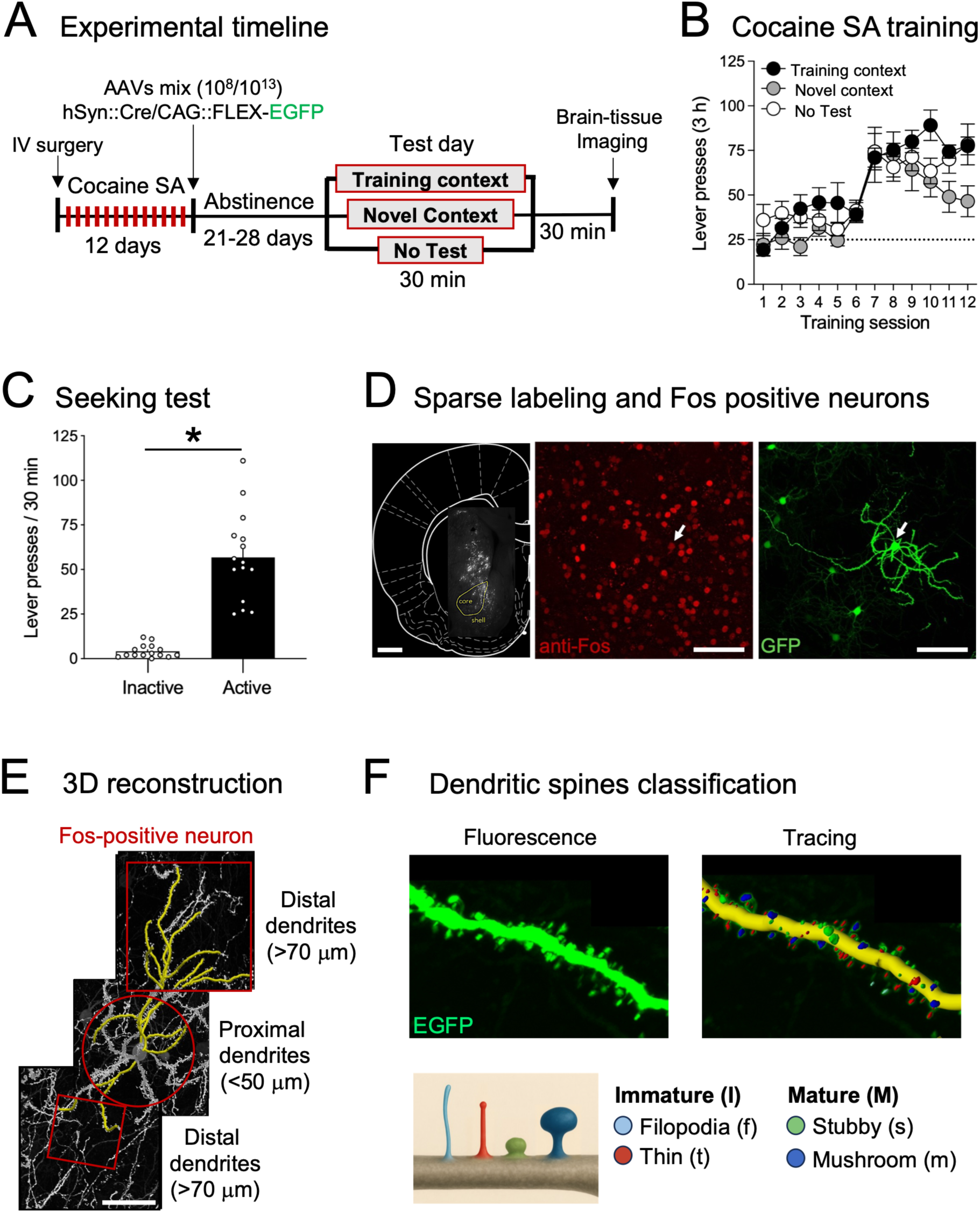
Experimental design, behavior and methodological strategy for analyzing spine morphology of Fos-positive and Fos-negative neurons in NAc core. (A) Experimental timeline. (B) Self-administration training: number of active lever presses over 12 days of 3 h daily sessions for the 3 groups: Training context (n = 15), Novel context (n = 13), No test (n = 12). (C) Cue-induced seeking test: total number of active lever presses during cue-induced cocaine seeking (n = 15). Data are shown as mean ± SEM, *p < 0.01. (D) Left: Representative brain section showing sparse EGFP-labeled neurons in the dorsal and ventral striatum (scale bar = 1 mm). Middle: Confocal image showing Fos immunopositive nuclei (red channel, scale bar = 100 µm). Right: confocal image of the selected Fos-positive neuron fully filled with EGFP fluorescence (green channel, scale bar = 100 µm). Arrow indicates a EGFP-positive neuron that express Fos nuclear protein (E) Fos-positive neuron with corresponding tracing of both dendrite and spines using Neurolucida 360 (scale bar = 50 µm). (F) Top images: magnified images of a segment from a distal dendrite from the example Fos-positive neuron shown in panel E labeled with EGFP. Bottom images: cartoon illustration of dendritic spine types arranged by increasing maturity from left to right. Immature spines, including filopodia (light blue) and thin (red) spines; and Mature spines, including stubby (green) and mushroom (dark blue) spines.

### Dendritic spine analysis in activated and non-activated neurons

To identify morphological changes of dendritic spines on activated medium spiny neurons, we injected an AAV expressing Cre-dependent EGFP combined with a low-titer 10,000-fold dilution of AAV expressing Cre recombinase into the dorsal medial striatum and NAc shell and core after the last day of cocaine self-administration (Fig. 1D). All brain sections were immunostained for Fos expression to identify activated neurons. The dual approach of Fos labeling along with sparse GFP labeling allowed us to identify nearby Fos-positive and Fos-negative neurons with clearly distinguishable non-overlapping dendrites. Images of proximal and distal dendrites on both Fos-positive and Fos-negative neurons were captured and processed for 3D reconstruction (Fig. 1E) and then spines were traced using Neurolucida 360 and classified into 4 categories based on their shape as filopodia, thin, stubby or mushroom spines (Fig. 1F). We restricted subsequent analyses to the NAc core subregion due to the demonstrated role of this subregion mediating cocaine cue-induced relapse in rats [27,28] and collapsed filopodia and thin spines together as ‘immature’ spines, since there were not enough filopodia to analyze separately. All results for proximal dendrites were placed in Supplemental figures since we did not detect any alterations on these segments; however all of these comparisons were hampered by very low numbers of spines on the proximal dendrites.

### Spine type classification and spine density

The composition of mature spines (stubby and mushroom combined) versus immature spines (thin and filopodia combined) were not significantly different between groups for both distal (Fig. 2A, Left) and proximal (Fig. S1A, Left) dendritic segments. The composition for the separate spine types (immature, stubby, mushroom) also did not differ significantly between groups for both distal (Fig. 2A, Right) and proximal (Fig. S1A, Right) dendritic segments. Spine densities were also not significantly different between groups for All spines (all spine types combined) or for separate immature, stubby or mushroom spine types on distal (Fig. 2B) and proximal (Fig. S1B) dendritic segments.

**Figure 2.**
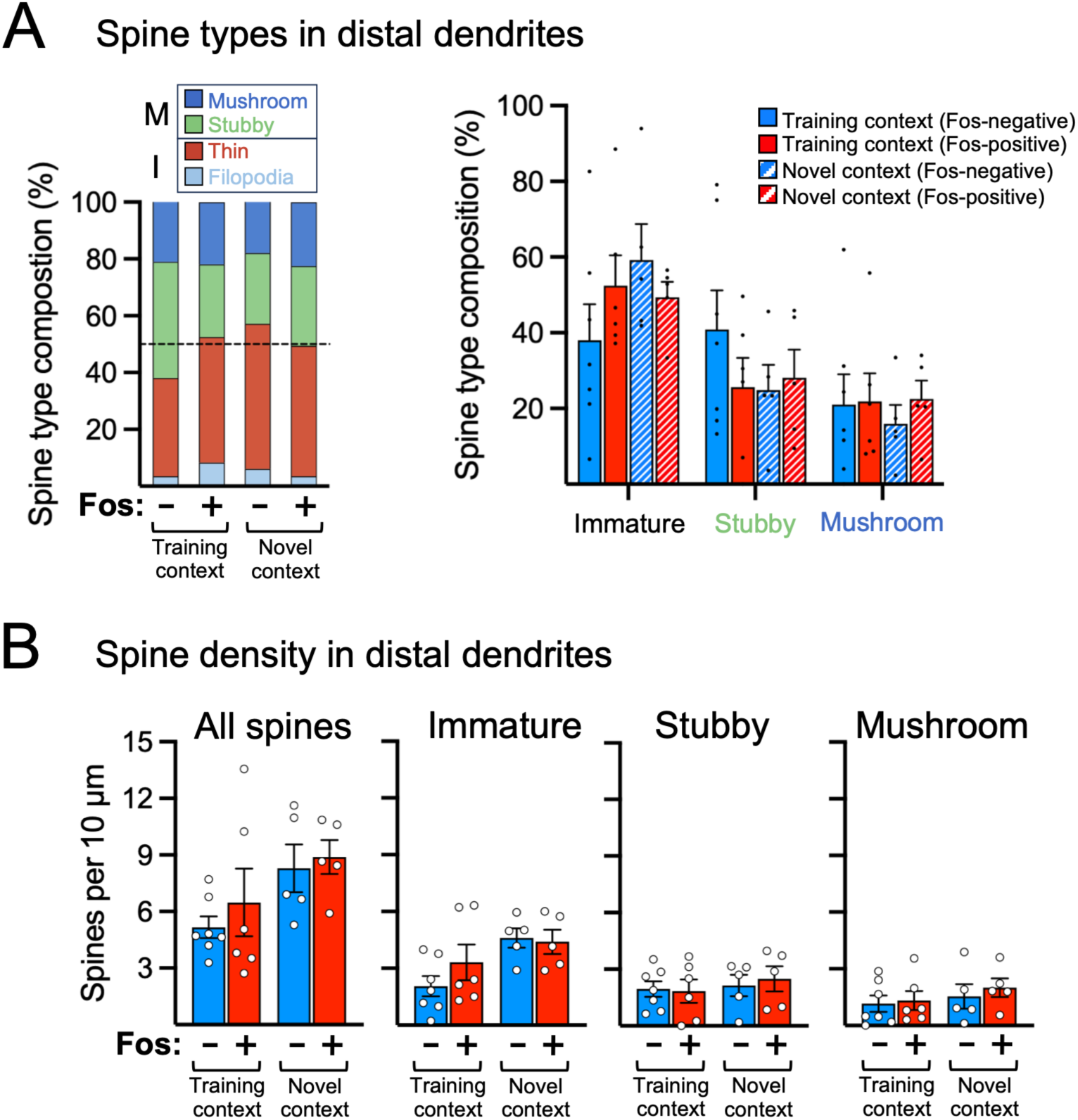
Dendritic spine type composition and spine densities on distal dendrites of NAc core Fos-positive (+) and Fos-negative (-) in the Training and Novel context groups. (A) Spine type composition: stacked bar graph (left side) showing the average percentage of each spine type: Immature (Filopodia, light blue; Thin, red), Stubby (green), and Mushroom (dark blue) spines in each group; bar graph (right side) showing percentage of each spine type in each experimental group. (B) Spine densities: number of spines per 10 µm for All spines (all spine types combined), Immature, Stubby, and Mushroom spines in each group. Data are shown as mean ± SEM; none of the groups were significantly different from each other (p > 0.05).

### Interspine intervals

Learning can alter the distribution of spines along the dendrite leading to functional effects associated with memory formation [12,18]. To analyze alterations of spine distribution, we quantified the distances between consecutive spines (interspine interval; ISI) in bins covering the range of 0-11.75 µm since previous studies demonstrated that spines can interact functionally when they are within 10-12 µm of each other [29,30]. For distal dendrites in the Training and Novel context groups, the cumulative distributions of ISI frequency were not significantly different between Fos-positive and Fos-negative neurons for All spines (Fig. 3A; all spine types combined), Immature spines (Fig. 3B), or Stubby spines (Fig. 3C). However, the cumulative distributions of ISI frequency for mushroom spines were significantly different between Fos-positive and Fos-negative neurons in the Training context group for the 2 µm bin that included 1.25-2.75 µm ISIs (*F*_7,56_ = 2.143, p = 0.0536), but not in the Novel context group (*F*_7,56_ = 0.711, p = 0.6630, (Fig. 3D). When we compared mushroom spines in only the 2 µm bin using 2-way ANOVA, the frequencies of spines in this bin were increased on Fos-positive versus Fos-negative neurons in the Training context group, but not in the Novel context group (p < 0.05, Fisher’s LSD test, Fig. 3D inset), indicating an increased proportion of spines found closer together on Fos-positive versus Fos-negative neurons. In proximal dendrites, the cumulative distributions of ISI frequencies were not different between Fos-positive and Fos-negative neurons for any spine types (Fig. S1C).

**Figure 3.**
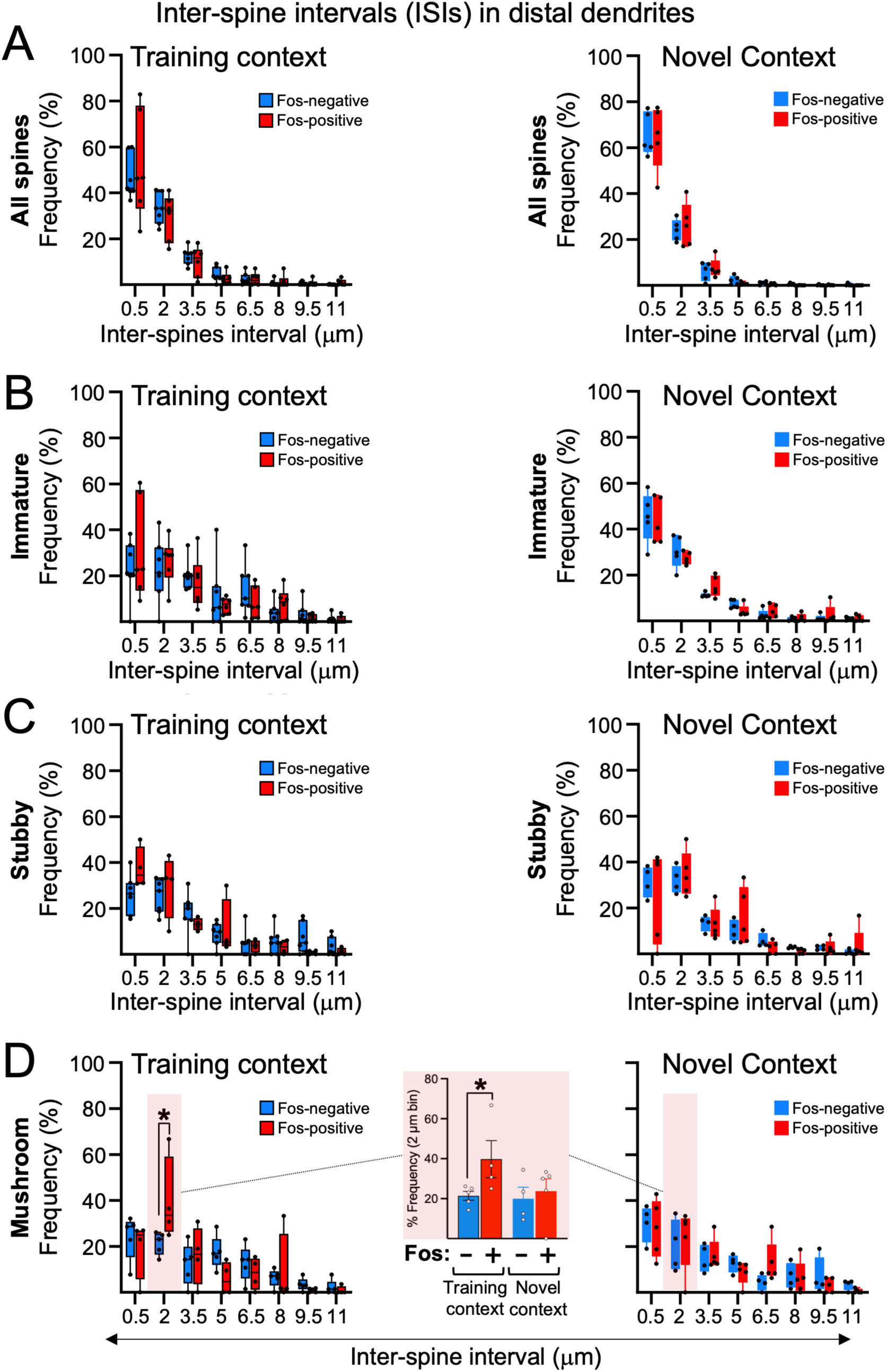
Distribution of spines expressed as interspine intervals (ISI) in distal dendrites of NAc core Fos-positive and Fos-negative neurons in the Training and Novel context groups. Box plots (min to max) representing the relative frequency (%) of ISIs, binned at 1.5 µm intervals (0.5 to 11 µm bin centers), for (A) All spines, (B) Immature, (C) Stubby and (D) Mushroom spines. Only mushroom spines in (D) showed a significant difference for ISIs between Fos-positive and Fos-negative neurons, and only for ISIs in the 2 µm bin (highlighted in pink). The bar graph in middle of panel D represents frequency of ISIs for only the 2 µm bins in the Training and Novel context groups; the frequency of spines in the 2 µm bin were significantly higher in Fos-positive (+) versus Fos-negative (-) neurons in the Training context group, but not in the Novel context group. Data are shown as mean ± SEM; *p < 0.05, n=4-5 neurons.

### Mushroom spine head diameters

We next investigated morphological alterations of mushroom spines due to their important role in learning and memory [14–16]. We started with head diameters within the larger category of All spine types. In distal dendrites, the cumulative distributions of head diameters were shifted to the left on Fos-positive (relative to Fos-negative) neurons in the Training context group (*F*_1,120_ = 36.98, p < 0.01, Fig. 4A), but not in the Novel context group (*F*_1,96_ = 0.004, p = 0.94), indicating decreased head diameters on Fos-positive neurons, specifically for spines in the 0.5 and 0.6 µm bins. However, 2-way ANOVA indicated group averages for head diameters were not significantly different between Fos-positive and Fos-negative neurons in the Training or Novel context groups (Fig. 4B). For mushroom spines in distal dendrites, the cumulative distributions of head diameters were also shifted to the left on Fos-positive (relative to Fos-negative) neurons in the Training context group (*F*_7,72_ = 2.703, p < 0.05), but not in the Novel context group (*F*_7,56_ = 0.346, p = 0.9288), indicating decreased head diameters on Fos-positive neurons, specifically for spines in the 0.5-0.8 µm bins (Fig. 4C). This time, 2-way ANOVA indicated group averages for head diameters were significantly different (*F*_1,16_ = 6.872, p < 0.05); subsequent posthoc analyses indicate head diameters were significantly decreased on Fos-positive (relative to Fos-negative) neurons in the Training context group (p < 0.05) but not in the Novel context group (Fig. 4D). Spine head diameters were also greater in the Training context (relative to Novel context) group for Fos-negative neurons (p <0.05), but not for Fos-positive neurons. In proximal dendrites, the cumulative distributions of head diameters were not different between Fos-positive and Fos-negative neurons for any spine type (Fig. S2A-D).

**Figure 4.**
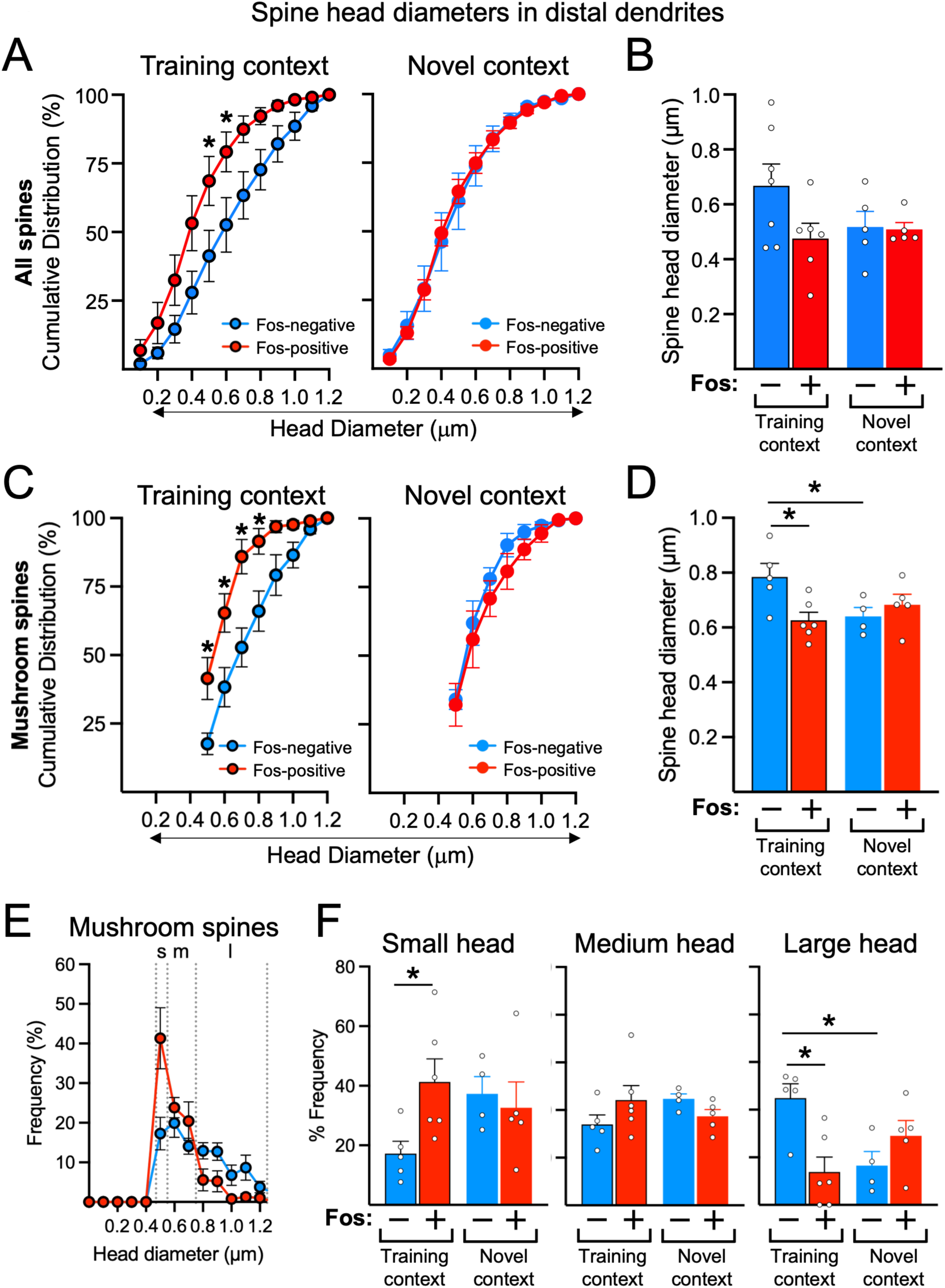
Spine head diameters on distal dendrites of NAc core Fos-positive (+) and Fos-negative (-) neurons in the Training and Novel context groups. Cumulative distribution of head diameters for (A) All spines (all spine types combined) and (C) mushroom spines. Bar graphs show the group averages of spine head diameters for (B) All spines and (D) mushroom spines. (E) Frequency distribution of mushroom spine head diameters divided into three components designated: Small (s), Medium (m) and Large (l) head mushroom spines. (F) Bar graphs show the average frequency of Small (0.47 to 0.54 µm), Medium (> 0.54 to 0.74 µm), and Large (> 0.74 µm) head mushroom spines across groups. Data are shown as mean ± SEM; *p < 0.05, n=4-6 neurons.

One of the two previous studies that examined spines on Fos-positive versus Fos neg neurons in drug-cue learning [23] found spine head diameters were increased on Fos-positive neurons, but not on Fos-negative neurons, 30 min following exposure to an amphetamine-paired context. Although they did not separate their head diameter data by spine types, they found increased numbers of spines with head diameters specifically in the 0.45-0.60 µm range but not for other spine head diameter ranges. To directly compare with their study, we reclassified our mushroom spines into three subcategories: Small (s; 0.47-0.54 µm), Medium (m; >0.54-0.74 µm), and Large (l; >0.74 µm) head diameters, based on three different components of the mushroom spine head diameter distribution (Fig. 4E). Following two-way ANOVA for the Small head subcategory (*F*_1,16_ = 4.045, p = 0.0615 for Fos x Context interaction), we found the frequency of mushroom spines was significantly increased on Fos-positive (relative to Fos-negative) neurons in the Training context group (p < 0.05), but not in the Novel context group (p= 0.8885) (Fig. 4F, Left). Within the Medium head subcategory, the frequencies of mushroom spines were not different between groups (Fig. 4F, Middle). Following two-way ANOVA for the Large head subcategory (*F*_1,16_ = 11.81, p < 0.01 for Fos x Context interaction), we found the frequency of mushroom spines was significantly decreased on Fos-positive (relative to Fos-negative) neurons in the Training context group (p <0.01), but not in the Novel context group (p = 0.3643) (Fig. 4F, Right). The frequencies of mushroom spines were also greater in the Training context (relative to Novel context) group for Fos-negative neurons, but not for Fos-positive neurons (p <0.05). In proximal dendrites, the frequencies of mushroom spines were not different between groups for the Small, Medium, and Large subcategories (Fig. S2E).

### Mushroom spine neck diameters

We examined neck diameters of mushroom spines because they affect how the spine heads and dendritic shafts are chemically and electrically compartmentalized, which have significant effects on spine function [31]. The cumulative distributions and group averages of neck diameters for all mushroom spines were not significantly different between Fos-positive and Fos-negative neurons in the Training and Novel context groups in distal (Fig. S3A,B) or proximal dendritic segments (Fig. S3C,D). However, when we analyzed neck diameters of mushroom spines in distal dendrites within the same Small, Medium and Large head subcategories described above, we found the cumulative distributions of neck diameters in the Small head group were shifted to the right on Fos-positive (relative to Fos-negative) neurons in the Training context group (*F*_11,108_ = 4.046, p < 0.01), but not in the Novel context group (*F*_11,84_ = 0.7002, p = 0.7350) (Fig. 5A), indicating increased neck diameters on Fos-positive neurons, specifically in the 0.1 and 0.2 µm bins (p < 0.01). We also calculated head/neck ratios separately for mushroom spines in the Small, Medium and Large head subcategories and found significantly lower ratios for only Small head mushroom spines on Fos-positive (relative to Fos-negative) neurons in the Training context group (p < 0.05), but not in the Novel context group (Fig. 5B). Cumulative distributions and head/neck ratios for mushroom spines in the Medium and Large head subcategories were not significantly different between Fos-positive and Fos-negative neurons in the Training or Novel context groups (Fig. 5C-F).

**Figure 5.**
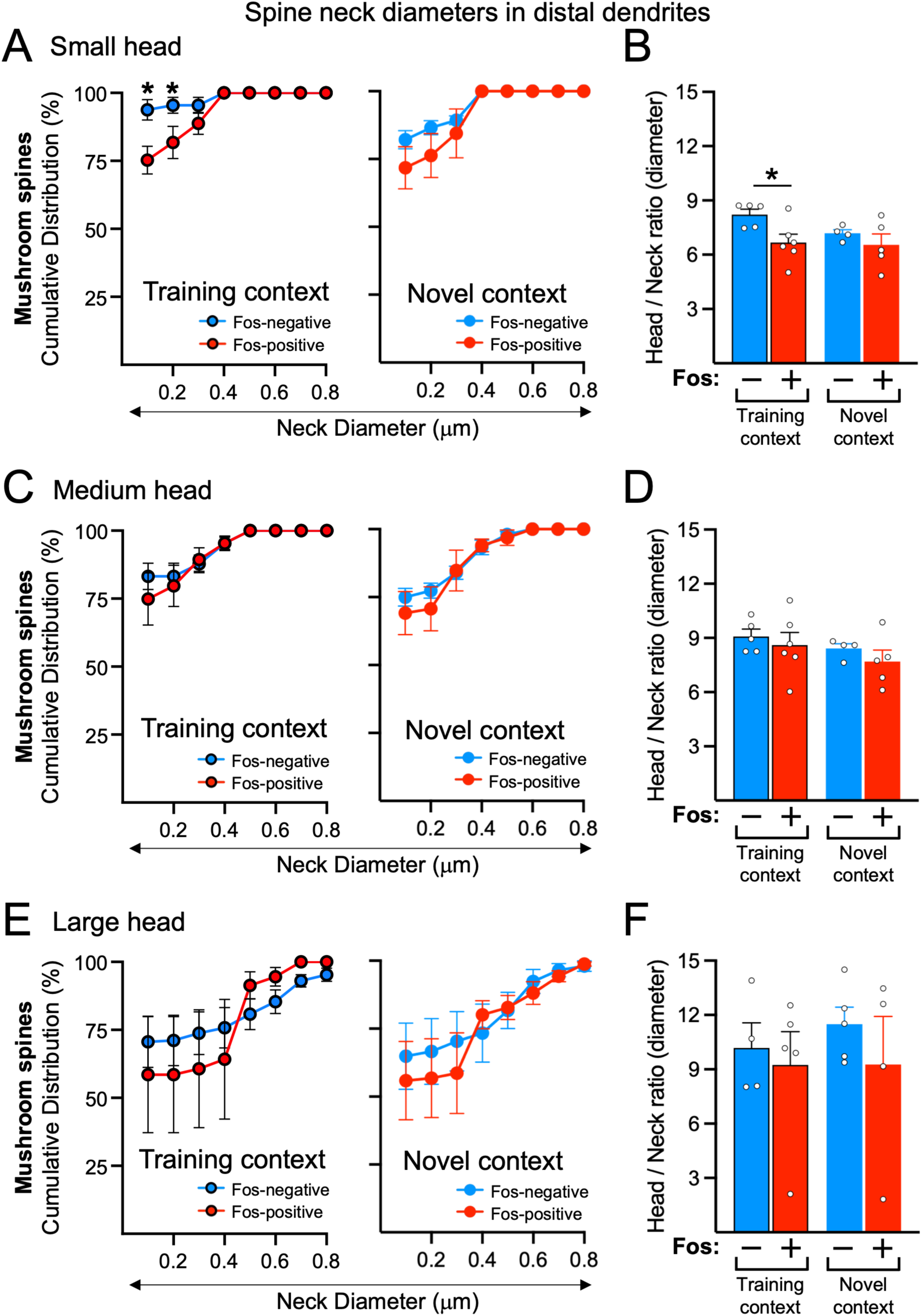
Spine neck diameters and head-to-neck ratios of mushroom spines in distal dendrites of NAc core Fos-positive (+) and Fos-negative (-) neurons in the Training and Novel context groups. Cumulative distribution of neck diameters for (A) Small, (C) Medium, and (E) Large mushroom spines. Bar graphs show the group averages of head-to-neck ratios for (B) Small, (D) Medium, and (F) Large mushroom spines across groups. Data are shown as mean ± SEM, *p < 0.05, n=4-6 neurons.

### Cocaine self-administration-specific spine alterations

Most studies of spine alterations in the drug learning literature examined spine alterations on distal dendritic segments or only secondary and higher order dendrites without distinguishing Fos-positive and Fos-negative neurons to assess the effects of drug self-administration training or repeated drug exposures [20,22,32–34]. To directly compare with these studies, we assessed the effects of cocaine self-administration alone by comparing spine alterations on distal dendritic segments in Naïve rats relative to those in rats that received the same cocaine self-administration training as the Training and Novel context groups but were kept in the home cage on test day (No Test group); since there was no acute activation of neurons in these two groups, nearly all neurons were Fos-negative. Spine types (Fig. S4A), spine densities (Fig. S4B) and ISIs (Fig. S5A-D) were not significantly different between Naïve and No Test groups. However, the cumulative distributions of head diameters were shifted to the left for the No Test (relative to the Naïve) group for All spines (specifically for the 0.5 µm and 0.6 µm bins) and Stubby spines (specifically for the 0.5-0.7 µm bins) (Fig. S6A), indicating decreased head diameters in the No Test group, while cumulative distributions for Mushroom spines were not significantly altered. However, the group averages for head diameters were not significantly different for any of these spine categories (Fig. S6B).

## Discussion

We found unique spine alterations on distal, but not proximal, dendrites of Fos-expressing (Fos-positive) neurons in nucleus accumbens (NAc) core that were activated during cocaine seeking. Specifically, mushroom spines on Fos-positive neurons had smaller head diameters and were found closer together within short 1.25-2.75 µm interspine intervals (ISIs) compared to those found on the surrounding Fos-negative neurons. Furthermore, these alterations were not found on Fos-positive neurons activated by exposure to a novel context instead of the training context, despite having identical cocaine self-administration histories. Altogether, these mushroom spine alterations describe a unique spine morphology signature for NAc core neurons selectively activated during cocaine relapse.

Although overall spine density was not altered, there was a large increase in the percentage of mushroom spines found clustered together on dendrites from Fos-positive, but not Fos-negative, neurons activated during relapse in the Training context group. Spine clustering is defined here as two or more spines within 5 µm of each other, based on previous studies [35]. In Fos-negative neurons, the ISI distribution decreased linearly from 20% in the 0.5 µm bin to almost 0% in the 11 µm bin. In contrast, the percentage of mushroom spines increased specifically in the 2 µm bin (1.25-2.75 µm) more than two-fold to almost half of all mushroom spines on Fos-positive neurons, indicating mushroom spines were clustered closer together on Fos-positive neurons rather than randomly dispersed along the dendrite, as it appears on Fos-negative neurons. Based on previous studies [30,35–38], we hypothesize that increased mushroom spine clustering on dendrites selectively on these Fos-positive neurons can enhance electrophysiological responsivity as well as increased synaptic plasticity contributing to storage and expression of the cocaine relapse memory.

Mushroom spine head diameters were smaller on Fos-positive neurons activated during cocaine relapse than on Fos-negative neurons. Based on previous studies [14,15], decreased mushroom spine head diameters in our study may reflect reduced postsynaptic density size, fewer glutamate receptors and lower electrophysiological responsivity; surprisingly, this would be expected to reduce synaptic efficacy of these Fos-positive neurons. Mushroom spine neck diameters were also wider (ranging 50-250 nm in the 0.1 and 0.2 µm bins) for the ‘Small head’ mushroom spines that were found more predominantly on Fos-positive neurons, leading to an overall reduction of head/neck ratio on these neurons. Based on previous studies [30,31], increased neck diameters can affect synaptic function by altering electrophysiological responses and the concentrations and time courses of signaling molecules (e.g. calcium) in the spine, which would also be expected to reduce synaptic efficacy of mushroom spines on Fos-positive neurons.

The lack of these spine alterations on Fos-positive neurons in the Novel context control group reveals a critical feature about the alterations on Fos-positive neurons in the Training context. In our previous studies, selective inactivation of Fos-expressing neurons that were previously activated by the cocaine-paired training context reduced subsequent relapse behavior, while inactivation of a different set of Fos-expressing neurons previously activated by a distinct set of stimuli in the novel context did not affect relapse behavior, even though the same number or more Fos-expressing neurons were activated by the novel context [3,4,39]. From these studies, we concluded that specific neuronal ensembles were recruited during training and then reactivated specifically by the drug training context to mediate relapse behavior. Since we used the same training context and novel context groups in the current study, we hypothesize that mushroom spine alterations on Fos-expressing neurons in the current training context group were produced on neuronal ensembles recruited during cocaine self-administration training and then observed only when the same neurons were reactivated and labeled for Fos expression in the same training context during cocaine relapse. In contrast, mushroom spine alterations were not observed on Fos-expressing neurons in the novel context group because Fos was expressed in a different set of neurons largely distinct from the training context neurons with the spine alterations. Note: if our mushroom spine changes were induced in neuronal ensembles activated during cocaine self-administration, one may expect to see similar mushroom spine alterations in the No test group that underwent cocaine self-administration but were kept in their home cages; however, it was not possible to identify and observe spine alterations on these neurons since Fos expression was not induced in this group.

Although numerous self-administration studies reported drug-induced spine alterations in the general neuronal population in several brain areas [20,22,40,41], only two studies examined spine alterations specifically on Fos-expressing neurons in NAc during recall of a drug memory [23,24]. Using a context-dependent amphetamine sensitization model and analyzing all spines together rather than separating into spine types, Singer et al found an increased number of medium-sized spines (0.45-0.60 µm head diameter) on Fos-expressing neurons activated by the drug-paired context, which is similar to the increased percentage of mushroom spines with ‘small head’ diameters in the same size range of 0.47-0.54 µm in our study (see Figure 5E). However, Singer et al did not find a similar decreased percentage of ‘large head’ mushroom spines as in our study but this may be due to them not separating into different spine types, particularly mushroom spines, prior to morphological analysis. In addition, Singer et al perfused their rat brains immediately after 30 min context exposure, while we allowed an additional 30 minutes in the home cage following the 30 min relapse test to permit more robust Fos expression. Since spine morphology can change rapidly within minutes following cocaine relapse and reinstatement [20], our mushroom spine head diameters may have decreased during the additional 30 min in the home cage.

Another recent study using Fos-TRAP2 mice also found dendritic spine morphological and electrophysiological alterations in NAc core neuronal ensembles during cue-induced cocaine reinstatement [24]. However, direct comparisons with our study are difficult because their mice underwent 10 days of extinction before ensemble labeling during reinstatement, followed by four additional extinction sessions before a second reinstatement test. Since extinction itself alters dendritic spines in NAc compared to acquisition [42,43], these experimental differences could change baseline spine properties on all MSNs prior to testing. Additionally, unlike our study, Flom et al did not analyze separate spine types, particularly mushroom spines. Future studies are needed to directly compare relapse-induced spine alterations on neuronal ensembles with and without prior extinction training.

Overall, the unique mushroom spine alterations we found on Fos-expressing neuronal ensembles activated during cocaine cue-induced relapse add to the growing number of molecular and cellular alterations found selectively on Fos-expressing ensembles. We propose these alterations constitute components of long-lasting engrams that encode drug-associated behaviors, with the current mushroom spine alterations representing components of a synaptic engram that physically encodes cocaine self-administration memories. To this end, spine redistribution or clustering is a slowly developing process during learning that takes days [35] and mushroom spines are long-lasting and known to play a role in memory storage [8]. Therefore, the redistribution of mushroom spines we observed following the relapse test is more likely a consequence of daily self-administration training than neural activity on test day, and capable of contributing to a long-lasting synaptic engram encoding the cocaine-associated memory. We are less certain of the stability of our observed head diameter alterations since they can change rapidly but these alterations are also capable of contributing to encoding memories [20,22]. Lastly, increased mushroom spine clustering and reduced head diameters found in our study would be expected to exert opposing effects on Fos-expressing neurons during cocaine relapse, enhancing and reducing synaptic responsiveness, respectively. Future studies are required to determine the time courses for development, stability and net functional effects of these spine alterations on neuronal function and their contributions to cocaine-associated learning and relapse.

## Methods (see Supplemental Extended Methods for details)

### Behavioral groups

All procedures followed the guidelines outlined in the *Guide for the Care and Use of Laboratory Animals (Ed 8;* http://grants.nih.gov/grants/olaw/Guide-for-the-Careand-Use-of-Laboratory-Animals.pdf). Forty male and female rats were trained 3 h/day to self-administer cocaine (0.75 mg/infusion days 1-6, 0.375 mg/infusion days 7-12) by pressing a lever paired with a cue light for 12 days followed by 21-28 days of forced abstinence (Experimental timeline in Fig. 1A). AAVs were injected one day after training. On test day, rats were separated into 3 groups that underwent either 30 min non-reinforced cocaine seeking in the training context (Training context group, n = 15), or 30 min exposure to a novel context (Novel context group, n= 13), or kept in their home cage (No test group, n =12). After the seeking test or novel context exposure, we returned rats to their home cages for 30 additional min before perfusing them. We added one additional control group of Naïve rats (n=7) that received IC surgeries, but not IV surgery or cocaine self-administration training, and were kept in the home cage for the duration of the experiment.

### Viral Injections

We injected a cocktail of 2 viruses: AAV1.hSyn.Cre.WPRE.hGH virus (diluted 10,000-fold to a final concentration 0.5 x 10^7^ vg/ml, catalog #105553-AAV1; Addgene) and AAV1-CAG-FLEX-EGFP-WPRE (final concentration 0.5 x 10^13^ vg/ml, catalog #51502-AAV1; Addgene) bilaterally into nucleus accumbens (NAc) shell, core and dorsal striatum (0.5 µl into each brain region) to visualize sparse EGFP-filled neurons.

### Fos immunohistochemistry and tissue clearing

We perfused rats with 4% paraformaldehyde and processed them for Fos immunohistochemistry using anti-Fos Ab (1:1000 dilution; catalog #5348, Cell Signaling). Tissue was cleared using the ScaleS procedure and coverslipped.

### Confocal imaging

We imaged proximal (<50 μm from the soma) and distal (>70 μm from the soma) dendritic segments from EGFP-labeled neurons using an Olympus FV3000 confocal microscope with a 30x silicone oil immersion objective lens (UPLSAPO30XS Super Apochromat, NA = 1.05, WD = 0.8 mm, Cat# N4212100, Evident). Only brain sections with well-filled EGFP-labeled individual neurons containing Fos-positive and/or Fos-negative neurons in the NAc region were selected for analysis: Training context group (n=12 total neurons from 7 brains), Novel context group (n=10 total neurons from 3 brains). No Test (n=5 neurons from 3 brains) and Naïve (n=5 neurons from 6 brains) groups were selected only for well-filled EGFP labeling. A detailed description of numbers of all rats, neurons, dendritic segments and spines counted are provided in Supplementary Table 1.

### Spine morphology analysis

We used Neurolucida 360 image software (MBF Bioscience) for 3D reconstruction and morphological analysis; we used previously defined morphological criteria to define spine types [44–46]. For each dendritic segment, we quantified spine density (spines per 10 µm), inter-spine interval, spine type composition, and individual spine morphology including head diameter and neck diameter.

### Statistical analyses

Behavioral data were analyzed using two-way ANOVAs (SPSS version 20 or GraphPad Prism v11 software) and Fisher’s PLSD post-hoc tests. *NOTE: For all spine analyses, we used each neuron as an independent sample (n-value). Cumulative distribution data were analyzed using two-way ANOVAs followed by Sidak’s correction for multiple posthoc comparisons of cumulative distributions of ISI, head diameter and neck diameter data. Group average data were analyzed using two-way ANOVAs followed by Sidak’s correction for multiple posthoc comparisons of head and neck diameter data and Fisher’s post-hoc test for ISI data.

## Author Contributions

FJR and BTH conceptualized the project, and designed and constructed the experimental setup and imaging strategy. FJR and RM optimized surgical and imaging protocols. FJR, RM, EU, DEO, EMH, AH, AK and LK performed behavioral and imaging experiments. FJR, RM, CMA and BTH developed analysis approaches. FJR and RM performed data analysis, generated figures and statistical outputs. FJR, RM, EU, CMA and BTH contributed critical intellectual input throughout the project. FJR, RM and BTH wrote the manuscript with feedback from all authors.

