## Supplemental Methods, Table, Figures for "Unique mushroom spine morphology signature in cocaine relapse ensembles in rat nucleus accumbens core"

**Supplemental files: Extended methods, Supplemental Figures and Legends**

**Supplemental extended methods**

*Subjects*

For the self-administration groups, we started with a total of 48 rats (25 male, 23 female) Sprague-Dawley rats (Charles River) weighing 250-300 g prior to IV surgery. We excluded 4 rats that did not learn to self-administer cocaine and 4 rats that died during the viral injection surgeries, leaving a total of 40 rats in the self-administration groups. A week before and after surgery, rats were housed individually under a reverse 12 h light/dark cycle (lights off at 8:00 AM). We included one additional control group of Naïve rats (n=7). Water was freely available in the rats' home cages, but food for cocaine-trained rats was restricted to 15 and 20 g of Purina rat chow for females and males, respectively, during all phases of the experiments. All procedures followed the guidelines outlined in the *Guide for the Care and Use of Laboratory Animals (Ed 8; http://grants.nih.gov/grants/olaw/Guide-for-the-Careand-Use-of-Laboratory-Animals.pdf ).*

*Intravenous surgeries*

We anesthetized rats with isoflurane (5% for induction; 2-3% for maintenance). We attached silastic catheters to a modified 22-gauge cannula cemented to a polypropylene mesh (Small Parts), then inserted the catheter into the jugular vein and fixed the mesh to the mid-scapular region of the rat as described previously (37-39). We injected the rats with 2.5 mg/kg of ketoprofen (Butler Schein) after surgery and the following day to relieve pain and decrease inflammation. Rats were allowed to recover for 10-11 d before starting cocaine self-administration training. During recovery and training, we flushed the catheters every day with 5 mg/ml of gentamicin (APP Pharmaceuticals) dissolved in sterile saline.

*Self-administration apparatus*

We trained and tested rats in Med Associates self-administration chambers located inside sound-attenuating cabinets. Each chamber had a red house light and two levers on the right side: a retractable active lever and a nonretractable inactive lever located 9 cm above the grid floor. Lever presses on the active retractable lever activated the infusion pump, whereas lever presses on the inactive nonretractable lever had no programmed consequences.

*Drugs*

Stock solutions of cocaine-hydrochloride in sterile saline were obtained from NIDA Pharmacy. (-)-cocaine-HCl was diluted in sterile saline for a final dose of 0.75 mg/kg/infusion for the first six days of training and a final dose of 0.375 mg/kg/infusion for days 7-12 of training.

*Behavioral procedures*

The experiment consisted of three phases: (1) self-administration training (3h/day for 12 d), (2) forced abstinence (21-28 days), and (3) cue-induced cocaine-seeking test (and novel context or home cage controls). The experimental timeline is shown in Fig. 1A.

*Phase 1: Cocaine self-administration (SA) training*

We trained rats to self-administer cocaine for 3-h/day for 12 d. The sessions began with extension of the active lever and illumination of the red house light, which remained on for the duration of the 3-h session. Active lever presses led to a 0.1 ml infusion of cocaine and 3.5 s of continuous tone-light cue located above the active lever. The maximum number of infusions was limited to 30 or 60 per session for days 1-6 or days 7-12, respectively. We trained the rats using a fixed-ratio-1 (FR-1) with a 20-s timeout reinforcement schedule in which rats did not receive a drug infusion after the active lever press. Inactive lever presses had no programmed consequences. All active and inactive lever presses were recorded during the duration of the session.

*Phase 2: Forced abstinence*

We housed rats in individual cages in the animal facility for 21-28 days. We gave rats ad libitum access to regular chow and water during this phase and handled them at least once per week.

*Phase 3: Cue-induced cocaine seeking test versus novel context and home cage controls*

Following abstinence phase, we separated the rats into three groups that underwent either 30 min non-reinforced cocaine seeking in the training context (Training context group, n = 15), or 30 min exposure to a novel context (Novel context group, n= 13), or kept in their home cage (No test group, n =12). For these three groups, rats were matched for previous cocaine-reinforced responding during the last 3 d of training. For the Training context group, the session started with 3.5 s of cocaine-cue presentation followed by extension of the active lever on the same FR-1, 20-s timeout reinforcement schedule but without drug. The inactive lever was always present during the test. After the seeking test (Training context group) or novel context exposure (Novel context group), we returned rats to their home cages for 30 additional min before perfusing them. Rats in the home cage group (No test group) were perfused immediately after removal from their home cages. We added one additional control group of Naïve rats (n=7) that received IC surgeries, but not IV surgery or cocaine self-administration training, and were kept in the home cage for the duration of the experiment.

*Viral Injections*

One day after training phase for the 3 cocaine-trained groups and for the 2 untrained groups, we injected a cocktail of two viruses: AAV1.hSyn.Cre.WPRE.hGH virus (diluted 10,000-fold to final concentration 0.5 x $10^7$ vg/ml, catalog #105553-AAV1; Addgene) and AAV1-CAG-FLEX-EGFP-WPRE (final concentration 0.5 x $10^{13}$ vg/ml, catalog #51502-AAV1; Addgene) bilaterally into the nucleus accumbens (NAc) shell, core and dorsal striatum after the last day of training to visualize sparse neurons filled with EGFP. We injected the virus first in the NAc shell and then the needle was raised to inject in the NAc core and again the needle was raised to inject in the dorsal striatum region. NAc shell coordinates were AP +1.4/1.5 mm (female/male), ML ±2.6 mm (10° angle), and DV -7.8 mm (0.5 µl per side); NAc core coordinates were AP +1.4/1.5 mm (female/male), ML ±2.6 mm (10° angle), and DV -7.1 mm (0.5 µl per side); dorsal striatum coordinates were AP +1.4/1.5 mm (female/male), ML ±2.6 mm (10° angle), and DV -5.0 mm (0.5 µl per side) and -4.5 mm (0.5 µl per side) with the nose bar set at -3.3 mm (40).

*Fos immunohistochemistry and tissue clearing*

We perfused rats from all 5 groups (3 cocaine-trained groups, 2 untrained groups) 60 min after the start of the stimulus (either Training context or Novel context) or directly from the home cage for the No test and Naïve groups. Rats were deeply anesthetized with isoflurane and perfused with 100 ml PBS followed by 400 ml of 4% paraformaldehyde in PBS. The brains were post-fixed in paraformaldehyde for 90 min and transferred to 30% sucrose in PBS solution at 4ºC for 2-3 d until they sank. Brains were frozen in powdered dry ice and kept at -80ºC until sectioning. Coronal sections (100 µm thick) were cut between bregma +2.28 mm and +0.36 mm for dorsal and ventral striatum (40). Free-floating sections were washed five times for 7 min each in PBS, incubated in in PBS containing 0.25% Triton X-100 (PBS-Tx) for 1 h, and incubated overnight at room temperature (22ºC) with anti-Fos antibody (1:1000 dilution; catalog #5348, Cell Signaling) in PBS-Tx for 3 nights or 72 h. Sections were washed again five times for 7 min each in PBS and incubated overnight or 16 h at 37ºC in Alexa Fluor 647 goat anti-rabbit secondary antibody (1:500 dilution; catalog #A21245, Invitrogen) in PBS-Tx. After washing five more times for 7 min each in PBS, sections were mounted onto Super Frost Plus slides (Fisher). The slides were dried at room temperature (22ºC) for 30 min and a hydrophobic pen (ImmEdger Hydrophobic Barrier Pen, Vector Laboratories) was used to make a physical barrier surrounding the brain sections. Sections were incubated at 37 ºC for 2 h in Scales SQ tissue clearing solution: 45 g D-sorbitol and 109.3 g urea were added to 200 ml distilled water, the solution was heated in a microwave and kept at 37 ºC. After removing the Scales SQ solution by tapping the slides several times onto a paper towel, the sections were incubated at room temperature for 2 h in Scales S4 solution: 80 g of D-sorbitol, 10 ml of glycerol, 48 g of urea and 20 ml of DMSO were added to 200 ml of distilled water, the solution was heated in a microwave and cooled to room temperature. The refractive index of Scales S4 solution allowed its use as a mounting medium for coverslipping.

*Confocal imaging and three-dimensional reconstructions*

We imaged the labeled neurons using an Olympus FV3000 confocal microscope, with the exception of one neuron imaged using a Leica Thunder microscope. Lasers (488 nm and 640 nm) were used to image endogenous EGFP and the AF647 signal for Fos immunoreactivity, respectively. Spines on proximal and distal dendritic segments of Fos-pos and Fos-neg neurons were imaged using a 30x silicone oil immersion objective lens (UPLSAPO30XS Super Apochromat, NA = 1.05, WD = 0.8 mm, Cat# N4212100, Evident), using a frame size of 1024 × 1024 pixels and a digital zoom of 3.5x to capture 121.218 x 121.218 μm per frame. Proximal (<50 μm from the soma) and distal (>70 μm from the soma) dendritic segments were scanned along the z-axis using 0.2 μm intervals (z-step size) and 0.8 μm Airy disk size. Final Z-stack images of the dendritic segments with spines were saved using Fluoview software (FV31S-SW, Olympus) for 3D reconstruction and morphological analysis.

*Neuron selection for imaging and spine analysis*

Only brain sections with well-filled EGFP-labeled individual neurons containing Fos-positive and/or Fos-negative neurons in the NAc region were selected for analysis: Training context (n=12 total neurons from 7 brains), Novel context group (n=10 total neurons from 3 brains), No test (n=5 neurons from 3 brains) and Naïve (n=5 neurons from 6 brains) groups were selected only for well-filled EGFP labeling since we can observe only Fos-neg neurons in these sections. A detailed description of numbers of all rats, neurons, dendritic segments and spines counted are provided in Supplementary Table 1.

**Supplementary Table 1**. Overall descriptive source of data from all analyzed spines

| Group | Fos IR | Distance from soma | Total rats | Total neurons | Total dendritic segments | Total dendritic length (μm) | Total all combined spines | Total filopodia spines | Total thin spines | Total stubby spines | Total mushroom spines |
|---|---|---|---|---|---|---|---|---|---|---|---|
| Training context | Fos-negative | distal | 6 | 7 | 39 | 1925 | 980 | 33 | 348 | 366 | 233 |
| | | proximal | 6 | 7 | 36 | 1093 | 387 | 16 | 171 | 139 | 61 |
| | Fos-positive | distal | 5 | 6 | 22 | 1317 | 841 | 40 | 393 | 232 | 176 |
| | | proximal | 4 | 5 | 37 | 1197 | 267 | 43 | 125 | 54 | 45 |
| Novel context | Fos-negative | distal | 3 | 5 | 43 | 2568 | 2315 | 100 | 1171 | 629 | 415 |
| | | proximal | 2 | 4 | 54 | 1437 | 1115 | 48 | 521 | 319 | 227 |
| | Fos-positive | distal | 3 | 5 | 32 | 1859 | 1772 | 43 | 790 | 573 | 366 |
| | | proximal | 2 | 4 | 39 | 1165 | 920 | 17 | 342 | 327 | 234 |
| No Test | Fos-negative | distal | 3 | 5 | 40 | 2180 | 2040 | 47 | 775 | 727 | 491 |
| | | proximal | 3 | 5 | 63 | 1476 | 819 | 38 | 304 | 331 | 146 |
| Naïve | Fos-negative | distal | 5 | 5 | 58 | 2632 | 2296 | 30 | 584 | 1110 | 572 |
| | | proximal | 5 | 5 | 87 | 2731 | 2247 | 44 | 720 | 843 | 640 |

*Morphological analysis of dendritic spines*

We used Neurolucida 360 image software (MBF Bioscience) for morphological analysis. For each neuron, multiple proximal and distal dendritic segments with lengths between 50-100 μm were analyzed within 1-3 regions of interest/frames (121.218 μm x 121.218 μm). Spine morphology data from different distal segments from the same neuron were combined. Dendrites were traced manually and refined using the smooth function, with diameter adjusted (1.6-3 μm) to match the actual dendrite width. Dendritic spines were detected and classified using Neurolucida 360. We used previously defined morphological criteria developed in previous fluorescence-based automated spine analyses (41-43) based on four morphological parameters: head-to-neck ratio, length-to-head ratio, mushroom head size, and filopodium length. Specifically, spines with a defined neck (head-to-neck ratio > 1.1) and head diameter ≥ 0.47μm were classified as mushroom. Spines with a defined neck but a smaller head were further classified based on length: those with a length-to-head ratio ≥ 2.5 and total length ≥ 3μm were classified as filopodia, while those with a length-to-head ratio ≥ 2.5 but length < 3μm were classified as thin. Spines lacking a defined neck (head-to-neck ratio < 1.1) and a length-to-head ratio <2.5 were classified as stubby. For each dendritic segment, we quantified spine density (spines per 10 μm), inter-spine interval, spine type composition (percentage of each spine type), and individual spine morphology including head diameter and neck diameter. For all spines, immature, stubby and mushroom spines, the distance between consecutive spines was measured from the base of the first spine to the base of the second spine.

*Statistical analyses*

Behavioral data were analyzed using two-way ANOVAs (SPSS version 20, GLM procedure or GraphPad Prism v11 software). We followed up on significant ANOVA effects using Fisher's PLSD post-hoc test. *NOTE: For all analyses of spine characteristics, we used each neuron as an independent sample (n-value), instead of the more common use of each spine as an independent sample in previous studies; on average, we assessed 1-3 neurons per rat (see *Neuron selection for imaging and spine analysis* section above for more detail). Cumulative distribution data for Fos-positive and Fos-negative neurons were analyzed using two-way ANOVAs with two within-subjects factors: Neuron type (Fos-positive, Fos-negative) and Bin number, calculated separately for the Training context and Novel context groups (GraphPad Prism v11 software). We followed up on significant ANOVA effects using Sidak's correction for multiple posthoc comparisons of cumulative distributions of ISI, head diameter and neck diameter data. Group average data for these data were analyzed using two-way ANOVAs with between-subjects factor Context (Training, Novel) and within-subjects factor Neuron type (Fos-positive, Fos-negative). We followed up on significant ANOVA effects using Sidak's correction for multiple posthoc comparisons of head and neck diameter data and Fisher's post-hoc test for ISI data.

**Supplemental figure legends**

**Figure S1**. Dendritic spine composition, spine densities, and distribution of spines expressed as interspine intervals (ISI) on proximal dendrites of NAc core Fos-positive (+) and Fos-negative (-) neurons in the Training and Novel context groups. (A) Spine type composition: stacked bar graph (left side) showing the average percentage of each spine type: Immature (Filopodia, light blue; Thin, red), Stubby (green), and Mushroom (dark blue) spines in each group; bar graph (right side) showing percentage of each spine type in each experimental group. (B) Spine densities: number of spines per 10 μm for All spines (all spine types combined), Immature, Stubby, and Mushroom spines in each group. (C) ISI frequencies: Box plots (min to max) representing the relative frequencies (%) of ISIs, binned at 1.5 μm intervals (0.5 to 11 μm bin centers) for All spines; numbers of spines on proximal dendrites were too few to analyze separate spine types. Data are shown as mean ± SEM; none of the groups were significantly different from each other ($p > 0.05$).

**Figure S2**. Spine head diameters on proximal dendrites of NAc core Fos-positive (+) and Fos-negative (-) neurons in the Training and Novel context groups. Cumulative distribution of head diameters for (A) All spines (all spine types combined) and (C) Mushroom spines. Bar graphs show the group averages of spine head diameters for (B) All spines and (D) Mushroom spines. (E) Bar graphs show the average frequencies of head diameters for Small (0.47 to 0.54 μm), Medium (> 0.54 to 0.74 μm), and Large (> 0.74 μm) mushroom spines across groups. Data are shown as mean ± SEM; none of the groups were significantly different from each other ($p > 0.05$).

**Figure S3**. Spine neck diameters of mushroom spines in distal and proximal dendrites of NAc core Fos-positive (+) and Fos-negative (-) neurons in the Training and Novel context groups. Cumulative distribution of neck diameters (from 0.1 to 0.8 μm) for all mushroom spines in (A) distal and (C) proximal dendrites. Bar graphs show the group average spine neck diameters for all mushroom spines in (B) distal and (D) proximal dendrites. Data are shown as mean ± SEM; none of the groups were significantly different from each other ($p > 0.05$).

**Figure S4**. Dendritic spine composition and spine densities on distal and proximal dendrites of NAc core Fos-negative neurons in Naïve (SA-) versus cocaine self-administration-No test (SA+) group. (A) Spine type composition: stacked bar graph (left side) showing the average percentage of each spine type: Immature (Filopodia, light blue; Thin, red), Stubby (green), and Mushroom (dark blue) in each group; bar graph (right side) showing percentage of each spine type in each experimental group. (B) Spine densities: number of spines per 10 μm for All spines (all spine types combined), Immature, Stubby, and Mushroom spines in Naive (SA-) group and cocaine self-administration-No test (SA+) group. Data are shown as mean ± SEM; none of the groups were significantly different from each other ($p > 0.05$).

**Figure S5**. Distribution of spines expressed as interspine intervals (ISI) on distal dendrites of NAc core Fos-negative neurons in Naïve (SA-) versus cocaine self-administration-No test (SA+) groups. Box plots (min to max) representing the relative frequency (%) of ISIs, binned at 1.5 μm intervals (0.5 to 11 μm bin center) for (A) All spines, (B) Immature, (C) Stubby and (D) Mushroom spines. Data are shown as mean ± SEM; none of the groups were significantly different from each other ($p > 0.05$).

**Figure S6**. Spine head diameters on distal dendrites of NAc core Fos-negative neurons in Naïve (SA-) versus cocaine self-administration-No test (SA+) groups. (A) Cumulative distribution of head diameters for (A) All spines (all spine types combined), Stubby and Mushroom spines. (B) Bar graphs show the group averages of spine head diameters for All spines, Stubby, and Mushroom spines. Data are shown as mean ± SEM; $^*p < 0.05$, n=4-7 neurons.

### Figure S1

Spine characteristics in proximal dendrites in Cocaine Cue versus Novel context rats

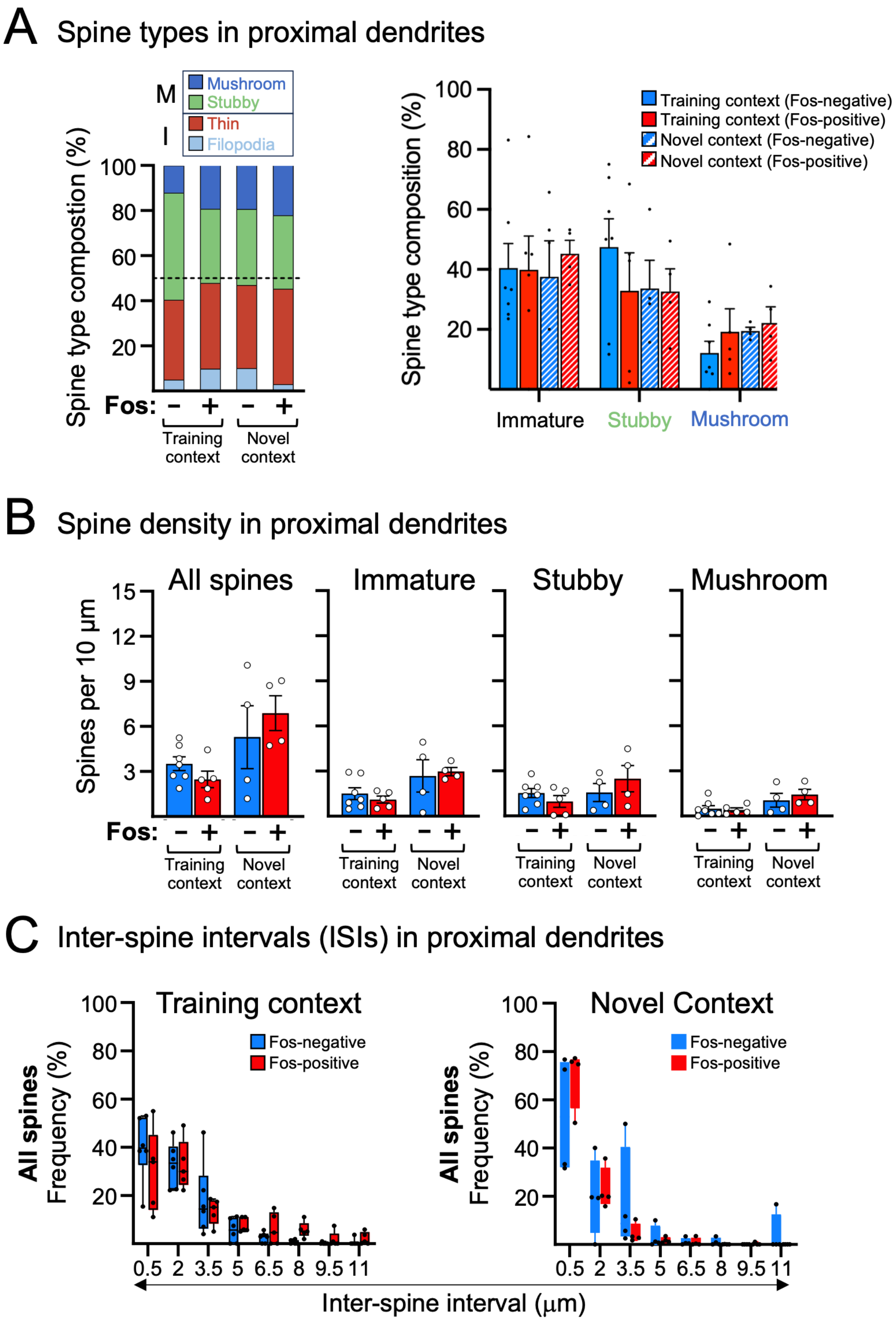

Figure S2 Mushroom spine head diameters in proximal dendrites in Cocaine Cue versus Novel context rats

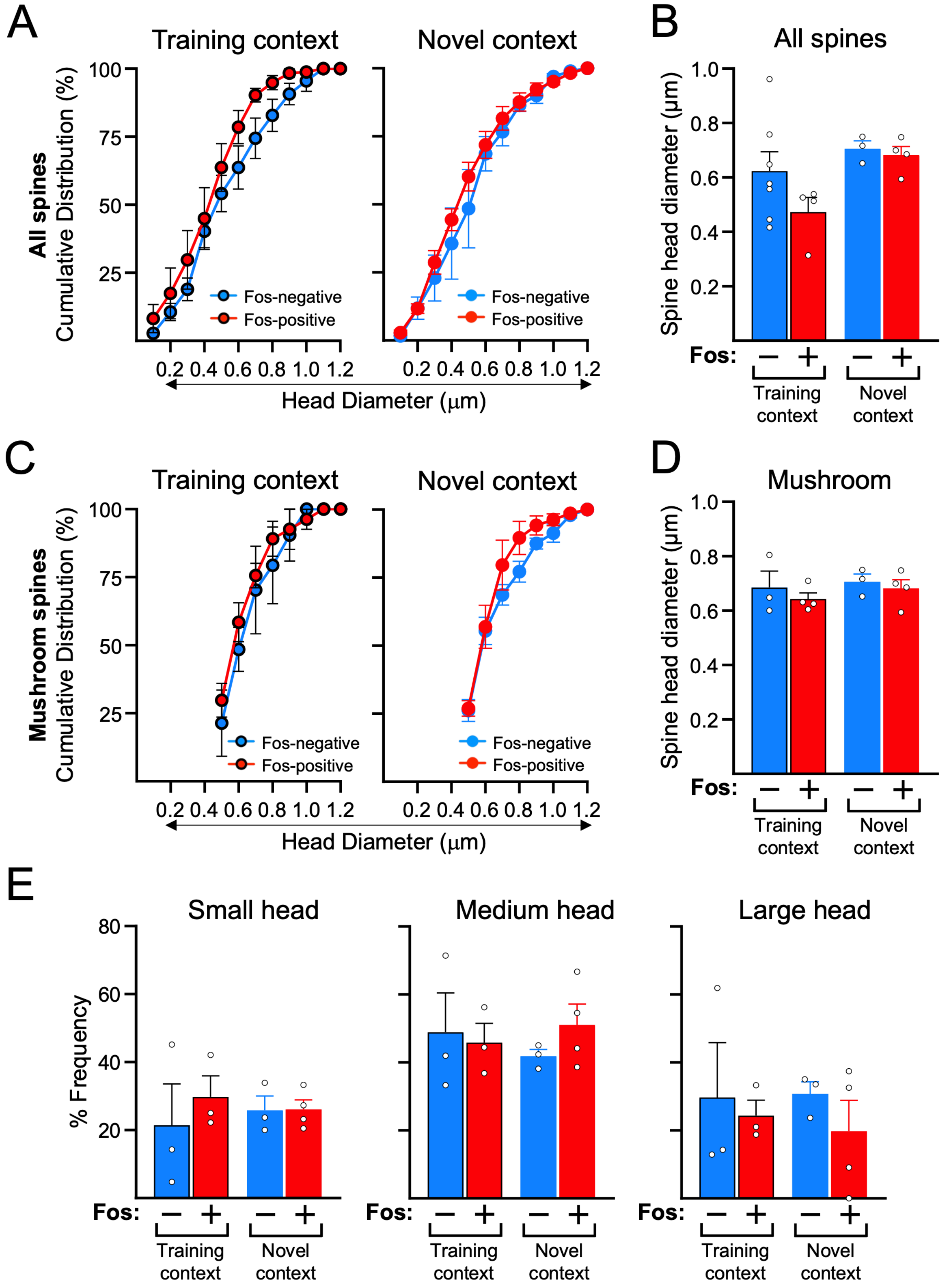

### Figure S3

Mushroom spine neck diameters in dendrites in
Cocaine Cue versus Novel context rats

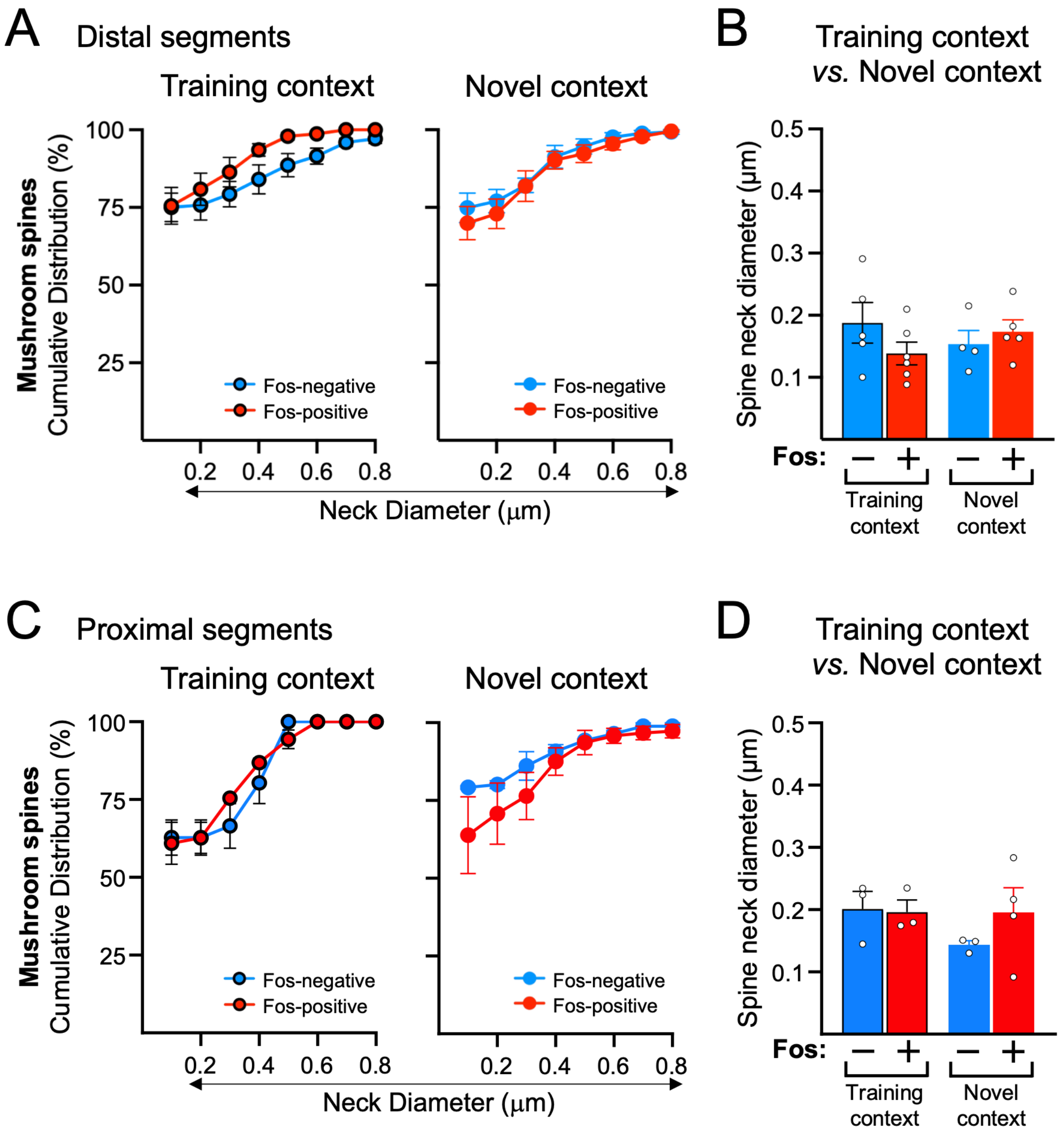

### Figure S4

Spine type and density for
Naïve versus Cocaine SA-trained rats

**A** Spine types in distal and proximal dendrites

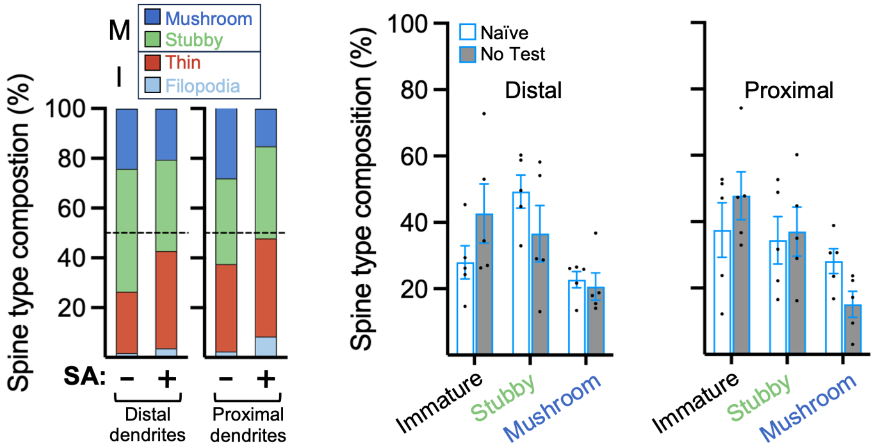


**B** Spine density in distal and proximal dendrites

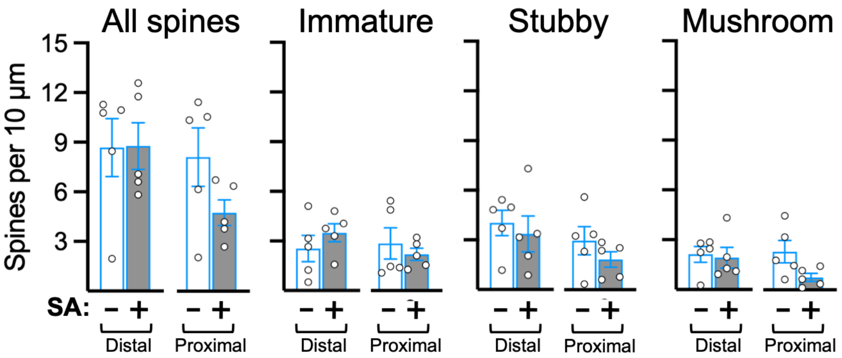

### Figure S5

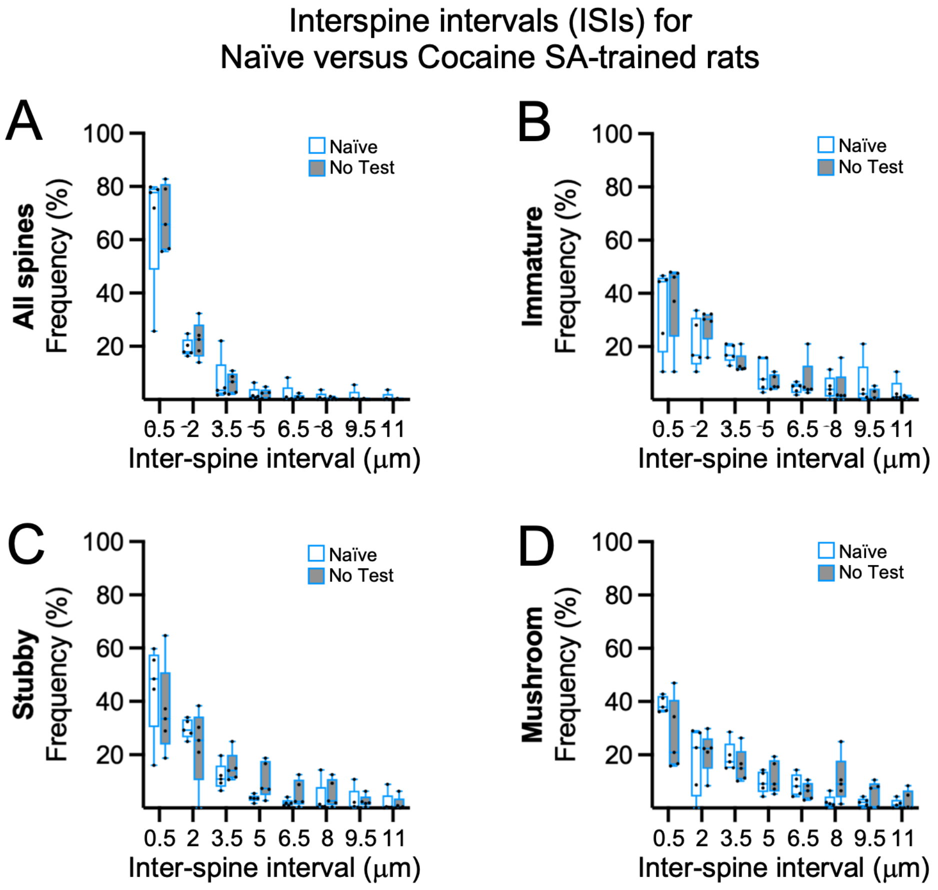

Interspine intervals (ISIs) for
Naïve versus Cocaine SA-trained rats
A
B
C
D
All spines
Immature
Stubby
Mushroom
Frequency (%)
100
80
60
40
20
Naïve
No Test
0.5
2
3.5
5
6.5
8
9.5
11
Inter-spine interval (μm)

Figure S6

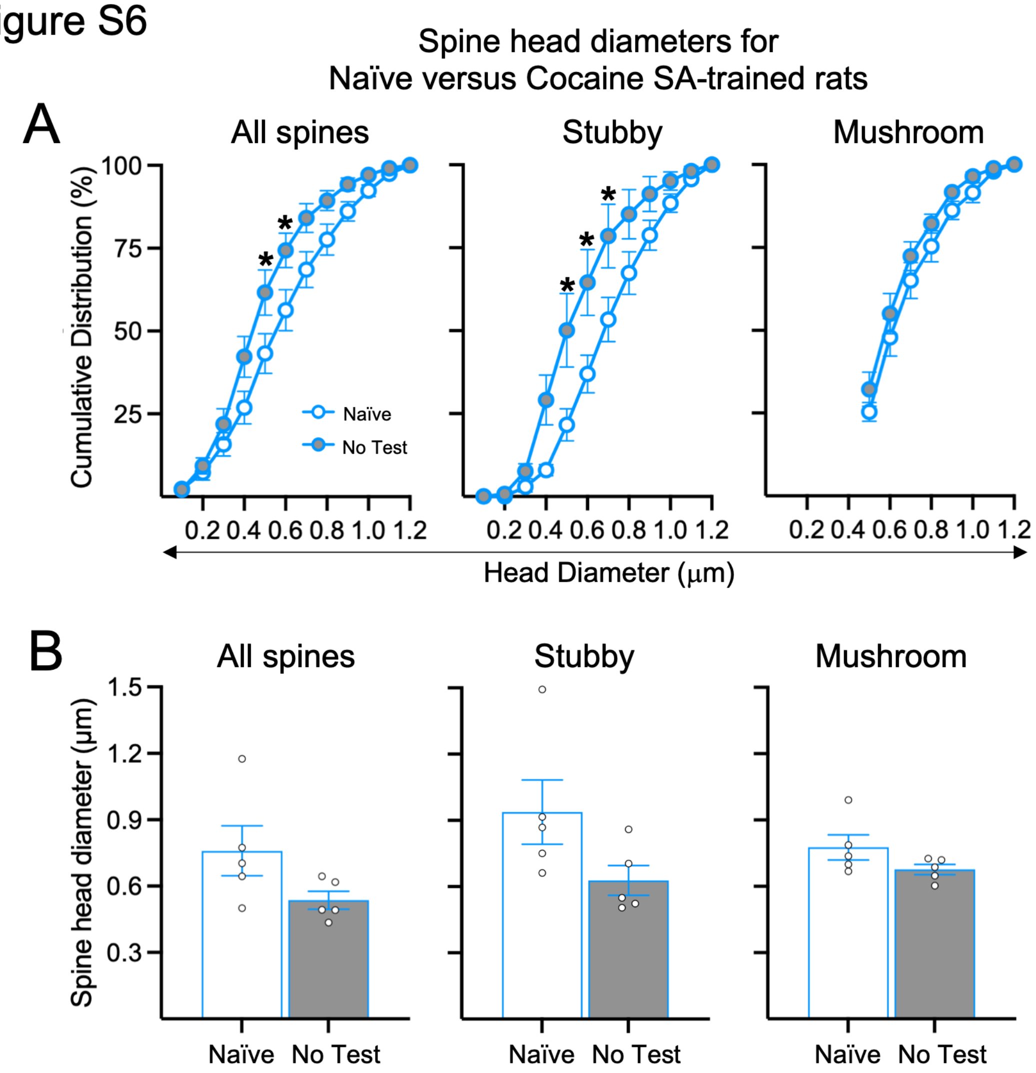

Spine head diameters for
Naïve versus Cocaine SA-trained rats
A
All spines
Stubby
Mushroom
Cumulative Distribution (%)
100
75
50
25
Naïve
No Test
0.2 0.4 0.6 0.8 1.0 1.2
Head Diameter (μm)
B
All spines
Stubby
Mushroom
Spine head diameter (μm)
1.5
1.2
0.9
0.6
0.3
Naïve
No Test
